# An *in vivo* reporter for tracking lipid droplet dynamics in transparent zebrafish

**DOI:** 10.1101/2020.11.09.375667

**Authors:** Dianne Lumaquin, Eleanor Johns, Joshua Weiss, Emily Montal, Olayinka Ooladipupo, Abderhman Abuhashem, Richard M. White

**Affiliations:** Cancer Biology and Genetics Program, Memorial Sloan Kettering Cancer Center, New York, NY, USA; Weill Cornell/Rockefeller/Sloan Kettering Tri-Institutional MD-PhD Program, Memorial Sloan Kettering Cancer Center, New York, NY, USA; Gerstner Sloan Kettering Graduate School of Biomedical Sciences, Memorial Sloan Kettering Cancer Center, New York, NY, USA; Developmental Biology Program, Memorial Sloan Kettering Cancer Center, New York, NY, USA

**Author notes:** These authors contributed equally to this work (Dianne Lumaquin, Eleanor Johns).

## Abstract

Lipid droplets are lipid storage organelles found in nearly all cell types from adipocytes to cancer cells. Although increasingly implicated in disease, current methods to study lipid droplets require fixation or static imaging which limits investigation of their rapid *in vivo* dynamics. To address this, we created a lipid droplet transgenic reporter in whole animals and cell culture by fusing tdTOMATO to Perilipin-2 (PLIN2), a lipid droplet structural protein. Expression of this transgene in transparent *casper* zebrafish enabled *in vivo* imaging of adipose depots responsive to nutrient deprivation and high-fat diet. Using this system, we tested novel regulators of lipolysis, revealing an unexpected role for nitric oxide in modulating adipocyte lipid droplets. Similarly, we expressed the PLIN2-tdTOMATO transgene in melanoma cells and found that the nitric oxide pathway also regulated lipid droplets in cancer. This model offers a tractable imaging platform to study lipid droplets across cell types and disease contexts.

## Introduction

Lipid droplets are cellular organelles which act as storage sites for neutral lipids and are key regulators of cellular metabolism (Farese & Walther, 2009). Lipid droplets are present in most cell types and are characterized by a monophospholipid membrane surrounding a hydrophobic lipid core (Olzmann & Carvalho, 2019). Cells maintain energetic homeostasis and membrane formation through the regulated incorporation and release of fatty acids and lipid species from the lipid droplet core (Jarc & Petan, 2019; Olzmann & Carvalho, 2019). Importantly, lipid droplets can assume various functions during cellular stress through the sequestration of potentially toxic lipids and misfolded proteins, maintenance of energy and redox homeostasis, regulation of fatty acid transfer to the mitochondria for β-oxidation, and the maintenance of ER membrane homeostasis (Olzmann & Carvalho, 2019; Petan et al., 2018). Moreover, recent work demonstrated that lipid droplets actively participate in the innate immune response (Bosch et al., 2020), and conversely, can be hijacked by infectious agents like hepatitis C virus to facilitate viral replication (Barba et al., 1997; Miyanari et al., 2007; Vieyres et al., 2020). The role of lipid droplets in metabolic homeostasis and cellular stress is critical across multiple cell types and has also been increasingly implicated in cancer (Petan et al., 2018). For example, lipid droplets can act as a storage pool in cancer cells after they take up lipids from extracellular sources, including adipocytes (Kuniyoshi et al., 2019; Nieman et al., 2011; Zhang et al., 2018).

While lipid droplets are ubiquitous across most cell types, they are essential to the function of adipocytes in regulating organismal energy homeostasis (Jarc & Petan, 2019). White adipocytes contain a large unilocular lipid droplet that is tightly regulated to mobilize fatty acids from the lipid droplet core (Heid et al., 2014; Zechner et al., 2017). Activation of lipolysis releases free fatty acids from the adipocyte lipid droplet which can be used by surrounding, non-adipose cell types to fuel energy production (Schoiswohl et al., 2010; Zimmermann et al., 2004). Dysregulation of lipid droplet function has been linked to a variety of pathophysiologies namely obesity (Olzmann & Carvalho, 2019). In addition, mutations in lipases required for lipolysis can lead to increased fat deposition and systemic metabolic abnormalities (Ahmadian et al., 2011; Haemmerle et al., 2006; Schoiswohl et al., 2010) in mouse models as well as the development of neutral lipid storage disease in humans (Fischer et al., 2007).

*In vivo* imaging of lipid droplets, either in adipocytes or in other cell types, is currently highly limited. Understanding these dynamics *in vivo*, rather than in fixed tissues, is important since the size of the lipid droplet can change very rapidly in response to fluctuating metabolic needs (Bosch et al., 2020; Fam et al., 2018). Much of adipose tissue imaging utilizes tissue fixation and sectioning, which can fail to preserve key aspects of the tissue structure (Berry et al., 2014; Xue et al., 2010). Whole mount imaging approaches in mice can be combined with adipocyte specific promoters, however, these methods still require tissue dissection and can be limited by tissue thickness (Berry & Rodeheffer, 2013; Chi et al., 2018).

Zebrafish offer a tractable model to address these limitations given the ease of high-throughput imaging of live animals. This is especially true with the availability of relatively transparent strains such as *casper*, which allow for detailed *in vivo* imaging without the need for fixation of the animal (White et al., 2008). Although less well studied than other vertebrates, zebrafish adipose tissue is highly similar to mammalian white adipose tissue and detailed work has classified the timing, dynamics, and location of zebrafish adipose tissue development (Minchin & Rawls, 2017). However, until now, the study of zebrafish adipose tissue has been limited to the use of lipophilic fluorescent dyes, which are restricted in their ability to read out dynamic changes over long periods of time (Fam et al., 2018).

Here, we report the development of an *in vivo* lipid droplet reporter using a *plin2-tdtomato* transgene in the *casper* strain. To date, transgenic lipid droplet reporters have been restricted to invertebrate model organisms such as *C. Elegans* and *Drosophila* (Kühnlein, 2011; Liu et al., 2014). We demonstrate that the reporter faithfully marks the lipid droplet which enables robust *in vivo* imaging. We show that this reporter can be applied to visualize adipocytes and to monitor adipose tissue remodeling in response to dietary and pharmacologic perturbations. Furthermore, we report the discovery of novel pharmacologic regulators of adipocyte lipolysis such as nitric oxide and demonstrate that several of these compounds can modulate adipose tissue area in our *in vivo* system. To facilitate the study of lipid droplets in novel contexts outside of adipocytes, we also generated a zebrafish melanoma cell line (ZMEL) (Heilmann et al., 2015) expressing *plin2-tdtomato* (ZMEL-LD). We confirm that this cell line can be used to monitor changes in lipid droplet production in response to both known and novel regulators of lipolysis. We anticipate that these models will be highly valuable as a high-throughput imaging platform to investigate lipid droplets in both adipose tissue biology as well as disease contexts such as cancer.

## Results

### An *in vivo* lipid droplet reporter using a PLIN2-tdTOMATO fusion transgene

To create a specific, fluorescent reporter for lipid droplets in zebrafish, we fused *tdtomato* to the 3’ end of the *plin2* cDNA. We chose *plin2* because it is a well-known lipid droplet associated protein that is ubiquitously expressed on lipid droplets across cell types (Olzmann & Carvalho, 2019). We generated stable transgenic zebrafish expressing *ubb:plin2-tdtomato* and sought to validate whether the construct faithfully marks lipid droplets (Figure 1A). White adipocytes are fat cells known for their large unilocular lipid droplet (T. Fujimoto & Parton, 2011; Heid et al., 2014) so we expected expression of the PLIN2-tdTOMATO fusion protein on the surface of the adipocyte lipid droplet (Figure 1A). Since the adipocyte lipid droplet occupies the majority of space in the cell (M. Fujimoto et al., 2020), existing methods to visualize zebrafish adipocytes rely on lipophilic dyes and lipid analogs which incorporate in the lipid droplet (Zhang et al., 2018). Thus in addition to labeling individual lipid droplets, we reasoned that the PLIN2-tdTOMATO fusion protein can also function as a reporter for adipocytes since these cells would have the largest and unilocular lipid droplets.

**Figure 1:**
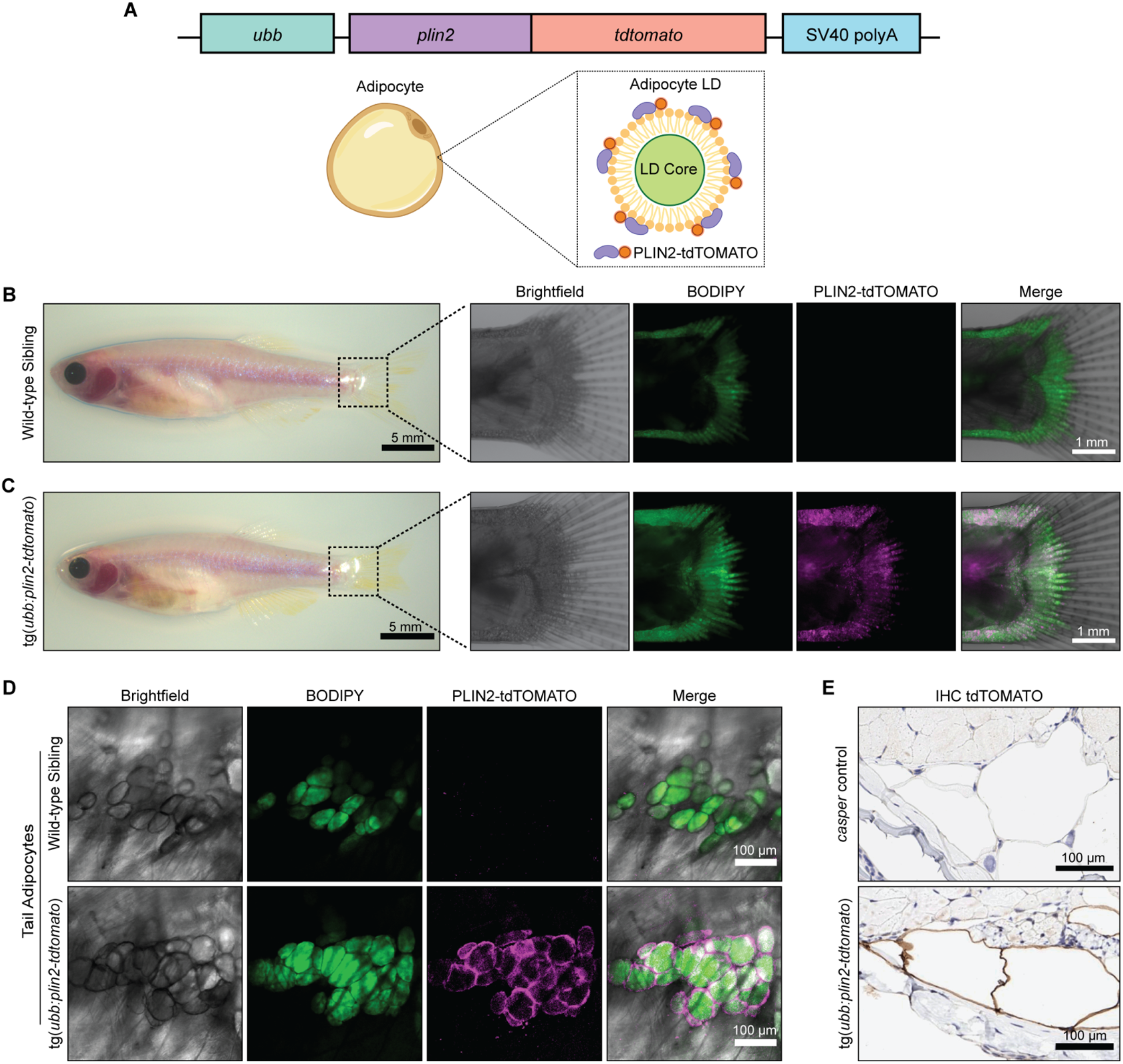
An *in vivo* lipid droplet reporter using a PLIN2-tdTOMATO fusion transgene. (A) Schematic of *ubb:plin2-tdtomato* construct injected into zebrafish and adipocyte lipid droplet labeled with PLIN2-tdTOMATO fusion protein. Widefield microscope images of adult (B) wild-type sibling and (C) tg(*ubb:plin2-tdtomato*) zebrafish. Box shows zoomed images of the fish tail with panels for brightfield, BODIPY, PLIN2-tdTOMATO, and merged. (D) Confocal images of fish tail adipocytes of adult wild-type sibling and tg(*ubb:plin2-tdtomato*) zebrafish. Panels show brightfield, BODIPY, PLIN2-tdTOMATO, and merge. (E) Adult *casper* and tg(*ubb:plin2-tdtomato*) zebrafish tails were fixed and immunohistochemistry conducted for tdTOMATO expression of tail adipocytes.

In adult zebrafish, subcutaneous adipocytes are known to reside proximally to the tail fin (Minchin & Rawls, 2017). When we imaged six month old adult *tg*(*ubb:plin2-tdtomato*) zebrafish, we detected PLIN2-tdTOMATO expression in the zebrafish tail fin adipocytes which colocalizes with BODIPY staining (Figure 1B, C). Lipophilic dyes such as BODIPY stain the lipid-rich core of the lipid droplet while lipid droplet resident proteins, such as PLIN2, localize to the lipid droplet membrane (Zhang et al., 2018). As expected, higher magnification images of tail adipocytes revealed that PLIN2-tdTOMATO expression was on the outside of the lipid droplet, whereas the BODIPY staining was on the interior of each droplet in the adipocyte (Figure 1D). Similarly, immunohistochemistry on the *tg*(*ubb:plin2-tdtomato*) zebrafish tail fin showed that adipocytes express tdTOMATO (Figure 1E). Taken together, this data demonstrates that the PLIN2-tdTOMATO fusion protein functions as a fluorescent lipid droplet reporter which can be applied to visualize adipocytes *in vivo*.

### The *tg*(*ubb:plin2-tdtomato*) is an *in vivo* reporter for visceral adipocytes

Visceral adipose tissue, otherwise known as abdominal fat, plays an important role in metabolism and participates in pathological processes of obesity, aging and metabolic syndromes (Tchernof & Després, 2013). Because PLIN2-tdTOMATO labeled subcutaneous adipocytes in the adult zebrafish tail fin, we wondered whether we could use the *tg*(*ubb:plin2-tdtomato*) zebrafish to visualize other adipose depots *in vivo* such as visceral adipocytes. In juvenile zebrafish at 21 days post-fertilization (dpf), visceral adipose tissue is composed of abdominal and pancreatic visceral adipocytes predominantly located on the right flank near the swim bladder (Figure 2A) (Minchin & Rawls, 2017). To determine whether *tg*(*ubb:plin2-tdtomato*) visceral adipocytes express PLIN2-tdTOMATO, we imaged around the swim bladder of juvenile zebrafish where we expect development of abdominal visceral adipocytes (Figure 2B). Visceral adipocytes visualized in brightfield co-stain for PLIN2-tdtomato and BODIPY, as we observed for subcutaneous adipocytes (Figure 2C). Immunohistochemistry of the juvenile *tg*(*ubb:plin2-tdtomato*) confirmed that the abdominal and visceral adipocytes express tdTOMATO (Figure 2D). Combined with the ability for high-throughput *in vivo* imaging in zebrafish, we sought to use *tg*(*ubb:plin2-tdtomato*) as a model to study lipid droplet dynamics in visceral adipocytes. One challenge we encountered was the auto-fluorescence from the zebrafish intestinal loops and gallbladder present in the tdTOMATO and GFP channels (Figure 2E). To remove background fluorescence, we developed an image analysis pipeline in MATLAB to segment the visceral adipocytes in the juvenile *tg*(*ubb:plin2-tdtomato*) (Figure 2E). Thus, *tg*(*ubb:plin2-tdtomato*) can be used as an *in vivo* model to visualize adipocytes with the benefits of avoiding staining steps and allowing for high-throughput image analysis in zebrafish.

**Figure 2:**
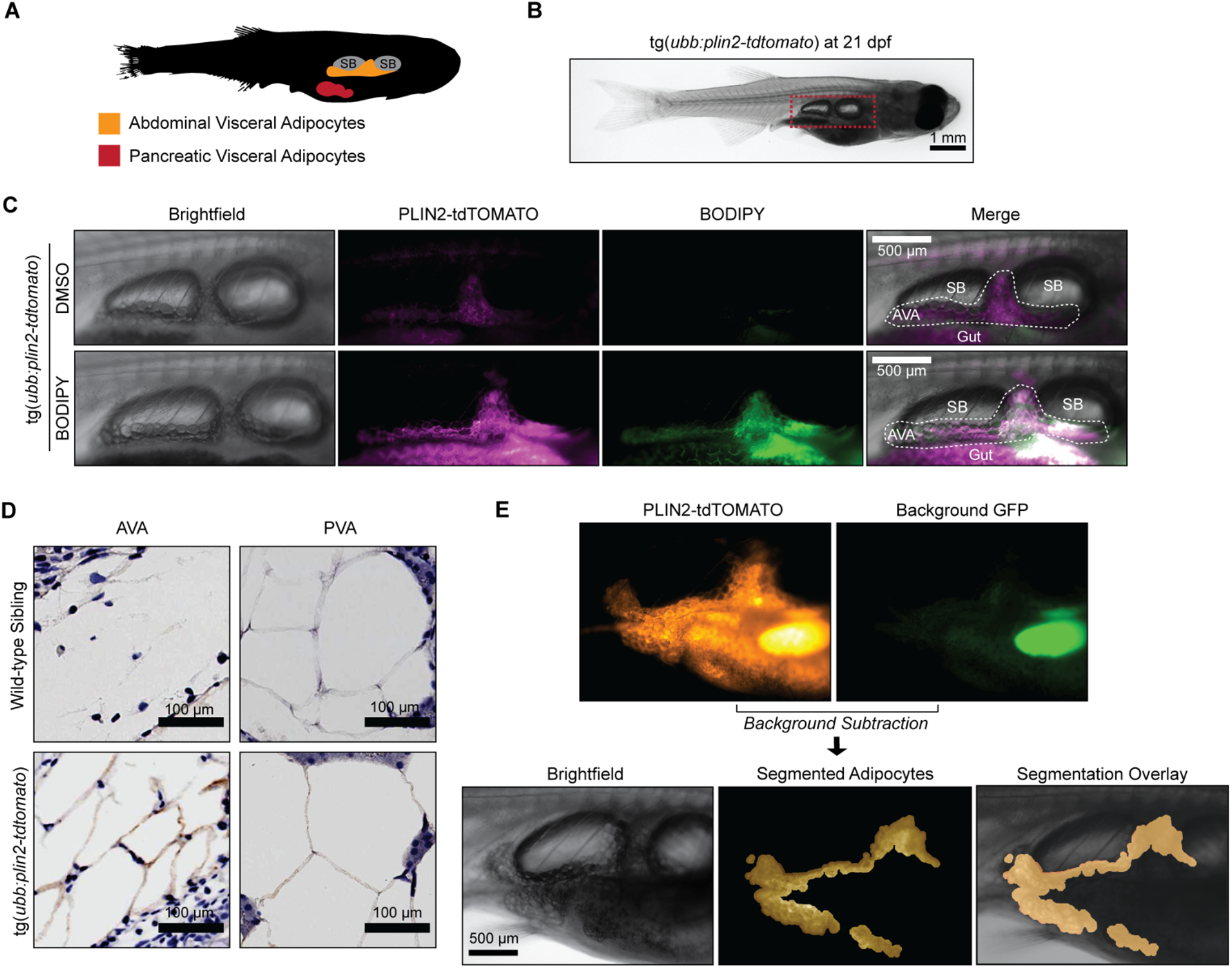
The tg(*ubb:plin2-tdtomato*) is an *in vivo* reporter for visceral adipocytes. (A) Schematic of visceral adipose tissue development in the juvenile zebrafish. Abdominal visceral adipocytes (orange) develop around the swim bladder (gray) and pancreatic visceral adipocytes (red) develop ventrally around the pancreas. (B) Brightfield image of juvenile tg(*ubb:plin2-tdtomato*) at 21 dpf. Red box indicates position of higher magnification images to visualize abdominal visceral adipocytes. (C) Widefield microscope images of juvenile *tg(ubb:plin2-tdtomato)* visceral adipocytes co-stained with DMSO or BODIPY. Panels show brightfield, PLIN2-tdTOMATO, BODIPY, and merge. Visceral adipocytes marked with white dash surrounding the swim bladder (SB) and gut. (D) Juvenile wild-type sibling and tg(*ubb:plin2-tdtomato*) zebrafish were fixed and immunohistochemistry conducted for tdTOMATO expression of abdominal (AVA) and pancreatic (PVA) visceral adipocytes. (E) Representative image of computational segmentation of juvenile tg(*ubb:plin2-tdtomato*) adipocytes. PLIN2-tdTOMATO was background subtracted with GFP fluorescence. Bottom panels show brightfield, segmented adipocytes, and segmentation overlaid on brightfield.

### Diet and pharmacologically induced reduction in visceral adipose tissue area

After confirming that we could image visceral adipose tissue in *tg*(*ubb:plin2-tdtomato*), we wanted to test whether this could be a tractable platform to image adipose tissue remodeling. We first verified whether we could use *tg*(*ubb:plin2-tdtomato*) to track reduction in visceral adiposity. Fasting is a well-known mechanism for reducing adiposity, since it will induce lipolysis and lead to a reduction in the size of the adipocyte lipid droplet (Henne et al., 2018; Longo & Mattson, 2014; Rambold et al., 2015; Tang et al., 2017). To test this, juvenile zebrafish were given control feed or fasted for 2.5 days then imaged to measure standard length and adipose tissue area (Figure 3A). As expected, we observed a reduction in the segmented adipocyte area in the fasted zebrafish (Figure 3B). Using our image analysis pipeline, we measured a significant reduction in adipose tissue area with an average of 0.39 ± 0.03 mm^2^ for fed fish and 0.21 ± 0.03 mm^2^ for fasted fish (Figure 3C). The control fed fish had a longer average standard length compared to the fasted fish (9.76 ± 0.18 mm vs 8.73 ± 0.17 mm) which we attribute to food restriction disrupting zebrafish development during this developmental window (Figure 3D). We saw a similar reduction in fasted fish when normalizing adipose tissue area to standard length, similar to a Body Mass Index (BMI) in mammals (control feed = 0.040 ± 0.003 area/SL and fasted = 0.024 ± 0.003 area/SL) (Figure 3E).

**Figure 3:**
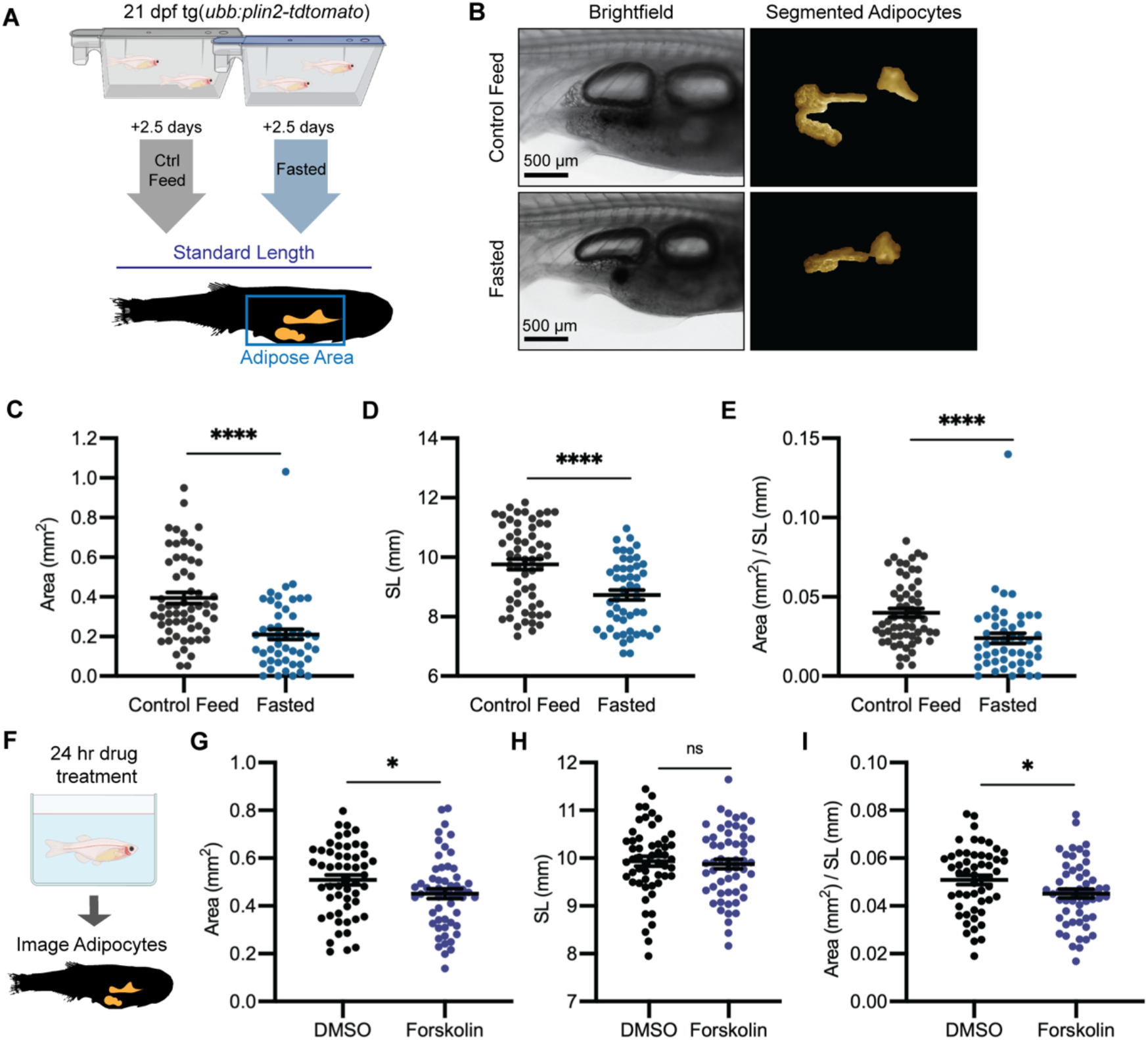
Diet and pharmacologically induced reduction in visceral adipose tissue area. (A) Schematic of experimental set-up for fasting experiment. 21 dpf *tg*(*ubb:plin2-tdtomato*) were fed control feed or fasted for 2.5 days and imaged to measure standard length and adipose area. (B) Representative images of zebrafish given control feed or fasted. Panels show images in brightfield and adipocyte segmentation. Image analysis pipeline resulted in measurements of adipose tissue (C) area, (D) standard length, and (E) area/standard length. Data points indicate individual fish for N = 4 independent experiments; Control feed n = 49; Fasted n = 59. Bars indicate mean and SEM. Significance calculated via Mann-Whitney test; **** p<0.0001. (F) Schematic of experimental set-up for Forskolin drug treatment. 21 dpf *tg*(*ubb:plin2-tdtomato*) were individually placed in 6 well plates with either DMSO or 5 μM Forskolin for 24 hours. Adipose tissue was imaged and analyzed for (G) area, (H) standard length, and (I) area/standard length. Data points indicate individual fish for N = 5 independent experiments; DMSO n = 53; Forskolin n = 55. Bars indicate mean and SEM. Significance calculated via Mann-Whitney test; * p<0.05.

In addition to fasting as a dietary perturbation, we also pharmacologically reduced adipose tissue. To achieve this, we used Forskolin, a drug which is known to induce lipolysis through cAMP signaling (Litosch et al., 1982). We treated juvenile zebrafish for 24 hours with either DMSO or 5 μM Forskolin and imaged the adipocytes (Figure 3F). We detected a reduction in both the adipose tissue area and normalized area to standard length in the Forskolin treated fish, but no differences in standard length (Figure 3G, H, I). Altogether, this data suggests that our PLIN2-tdTOMATO reporter faithfully reads out changes in the size of adipose tissue due to its capacity to sensitively detect lipolysis of the large lipid droplet in this tissue.

### High-fat diet leads to specific enlargement of visceral adipose tissue

Having shown that we could use *tg*(*ubb:plin2-tdtomato*) to image and measure reduction in adipose tissue, we tested whether we can use our model to detect an increase in adiposity. Zebrafish have been used as a model for diet-induced obesity and share pathophysiological perturbations seen in mammals, but few studies have focused on architectural changes of visceral adipose tissue (Chu et al., 2012; Landgraf et al., 2017; Oka et al., 2010). We sought to determine if we could detect increases in visceral adiposity from a high fat diet (HFD). We fed juvenile zebrafish with either control feed (12% crude fat) or HFD (23% crude fat) for 7 days and subsequently imaged the adipose tissue (Figure 4A, B). Remarkably after a week of HFD feeding, we observed that HFD fed fish developed notably increased visceral adiposity compared to the fish fed with control feed (Figure 4C). Quantification of the adipose tissue revealed that HFD led to an increase in adipose tissue area (control feed 0.42 ± 0.03 mm^2^ and HFD = 0.62 ± 0.03 mm^2^) and normalized area to standard length (control feed = 0.040 ± 0.002 area/SL and HFD = 0.061 ± 0.002 area/SL) (Figure 4D, F). Interestingly, we did not detect differences in the standard length of the fish (control feed = 10.09 ± 0.11 mm and HFD = 10.13 ± 0.11 mm), suggesting that this formulation of HFD leads to specific enlargement of visceral adipose tissue (Figure 4E). Our results demonstrate that *tg*(*ubb:plin2-tdtomato*) is an effective and unique tool to visualize visceral adipose tissue remodeling induced by HFD which can be widely applied to study obesity.

**Figure 4:**
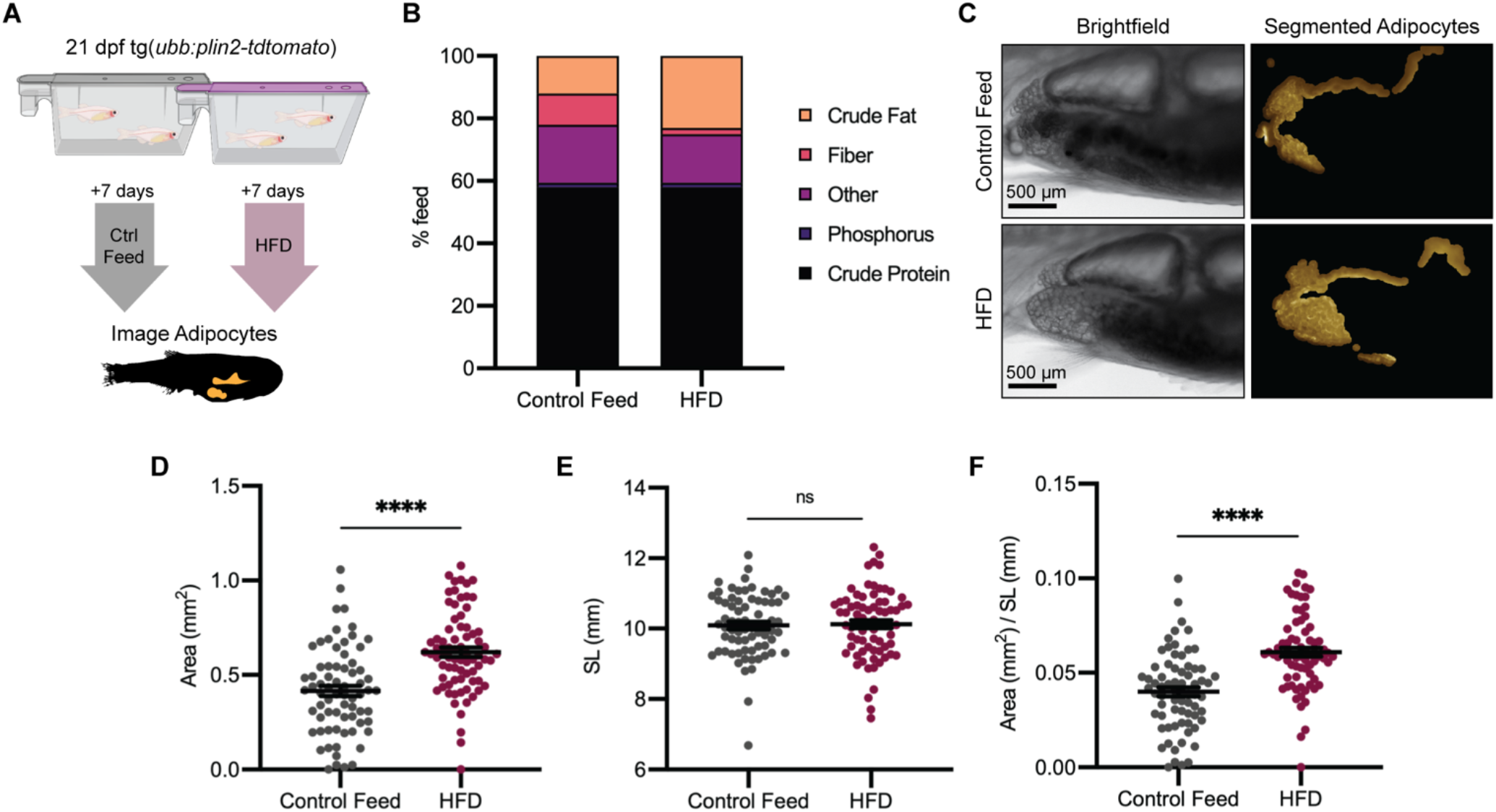
High-fat diet leads to specific enlargement of visceral adipose tissue. (A) Schematic of experimental set-up for high-fat diet (HFD) experiment. 21 dpf *tg*(*ubb:plin2-tdtomato*) were fed control feed or HFD for 7 days and imaged to measure standard length and adipose area. (B) Percent breakdown of nutritional content for control feed and HFD. (C) Representative images of tg(*ubb:plin2-tdtomato*) fed either control feed or HFD. Panels show images in brightfield and adipocyte segmentation. Image analysis pipeline resulted in measurements of adipose tissue (D) area, (E) standard length, and (F) area/standard length. Data points indicate individual fish for N = 3 independent experiments; Control feed n = 70; HFD n = 74. Bars indicate mean and SEM. Significance calculated via Mann-Whitney test; **** p<0.0001.

### A screen to discover novel compounds that modulate lipolysis and lipid droplets *in vivo*

To meet fluctuating nutritional needs of the cell, lipid droplets are remodeled through lipolysis to regulate lipid mobilization and metabolic homeostasis (Krahmer et al., 2013; Olzmann & Carvalho, 2019; Paar et al., 2012). As a major lipid depot for the body, white adipose tissue is critical to lipid availability and cycles through lipolytic flux in response to energy demands (Duncan et al., 2007). In disease contexts such as cancer, adipocytes undergoing lipolysis act as a lipid source for neighboring cancer cells (Lengyel et al., 2018). Adipocyte-derived lipids have been directly shown to promote cancer progression in ovarian (Nieman et al., 2011), breast (Balaban et al., 2017), and melanoma cancer cells (Zhang et al., 2018). Due to growing evidence of adipocyte and cancer cell cross-talk as a metabolic adaptation for tumor progression, there is significant interest in disrupting lipid transfer between adipocytes and cancer cells.

Leveraging our model to visualize lipid droplets in adipocytes, we became interested in identifying novel compounds that remodel adipocyte lipid droplets through lipolysis. In mammalian systems, the most commonly used cell line to study lipolysis are 3T3-L1 cells, which can be differentiated *in vitro* to resemble adipocytes (Zebisch et al., 2012). We first used the 3T3-L1 system to rapidly identify lipolysis inhibitors at high-throughput, and then test those hits using our zebrafish lipid droplet reporter. We reasoned that compounds which inhibit lipolysis *in vitro* would cause an increase in the size of the lipid droplets *in vivo*. To achieve this, we differentiated mouse 3T3-L1 fibroblast cells into adipocytes and conducted a chemical screen for compounds that inhibit lipolysis (Figure 5A), measured by quantifying glycerol in the media, a gold standard readout of lipolysis in this system (Hellmér et al., 1989). As a positive control, we used Atglistatin, an inhibitor of adipose triglyceride lipase (ATGL) which is known to be the rate limiting step of lipolysis and has been shown to inhibit lipolysis in cell lines and mouse models (Mayer et al., 2013; Schweiger et al., 2017). We confirmed that Atglistatin potently inhibits lipolysis in 3T3-L1 adipocytes (Figure 5B). We then screened through a library of 1,280 compounds of diverse chemical structures to find novel inhibitors of lipolysis. Overall, we found 29 out of 1,280 compounds which led to at least a 40% reduction in lipolysis as measured by glycerol release into the media. Looking more closely at the top 10 hits from this screen, we noted that 2 of the top 10 top hits (Auranofin and JS-K), both modulated nitric oxide (Figure 5A).

**Figure 5:**
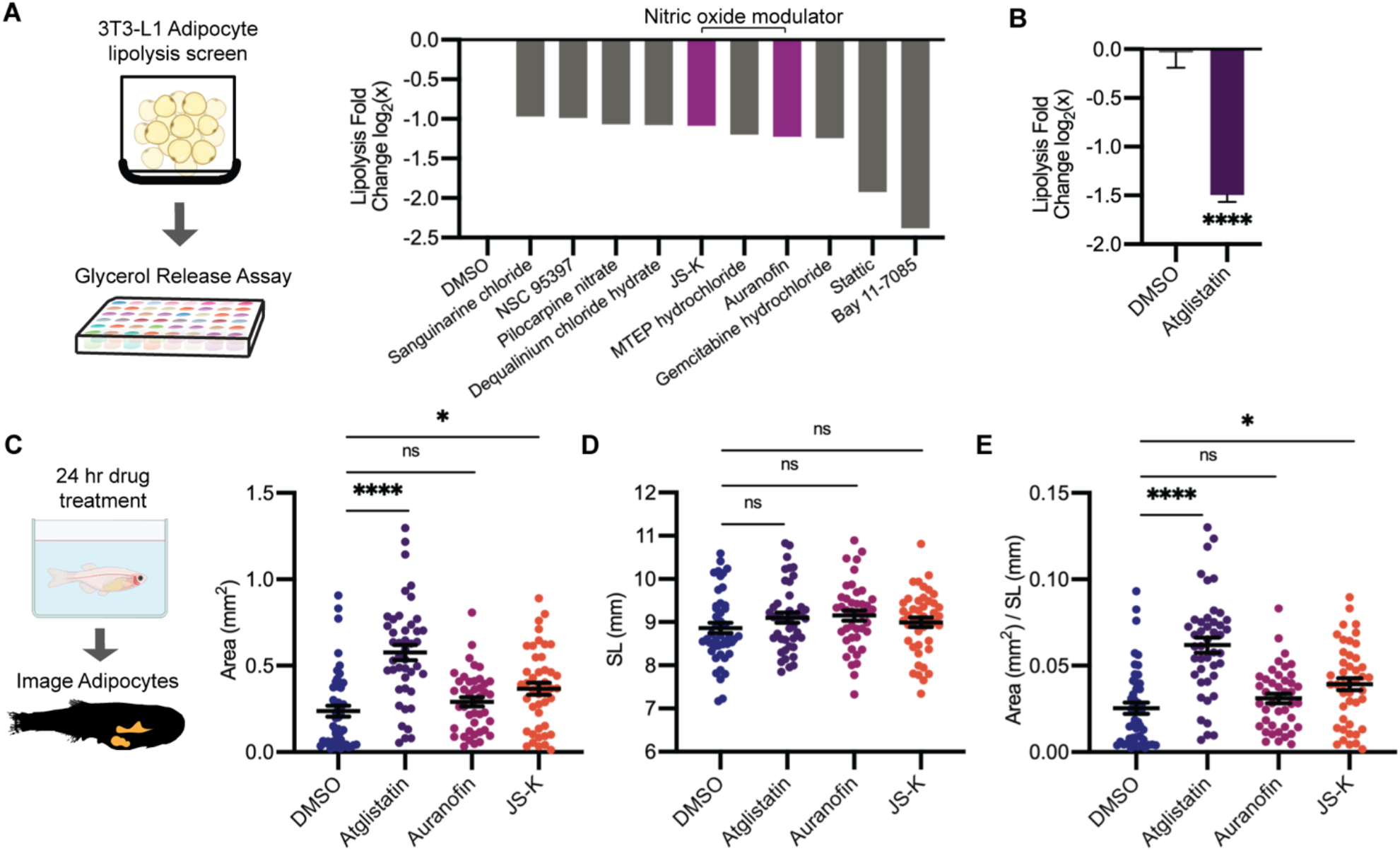
A screen to discover novel compounds that modulate lipolysis and lipid droplets *in vivo*. (A) Schematic of pharmacologic lipolysis screen in 3T3-L1 adipocytes using a glycerol release assay. Normalized values log2 transformed values for top ten drugs that inhibit lipolysis shown. Magenta indicates compounds that modulate nitric oxide. (B) Normalized values log2 transformed values for lipolysis inhibition in 3T3-L1 adipocytes using either DMSO or 100 μM Atglistatin. N = 5 independent experiments. Bars indicate mean and SEM. Significance calculated via unpaired t-test; ****p<0.0001. (C) Schematic of experimental set-up for drug treatment. 21 dpf *tg*(*ubb:plin2-tdtomato*) were individually placed in 6 well plates with either DMSO, 40 μM Atglistatin, 1 μM Auranofin, 1 μM JS-K for 24 hours. Adipose tissue was imaged and analyzed for (C) area, (D) standard length, and (E) area/standard length. Data points indicate individual fish for N = 4 independent experiments; DMSO n = 47; Atglistatin n = 44; Auranofin n = 42; JS-K n = 44. Bars indicate mean and SEM. Significance calculated via Kruskal-Wallis test; * p<0.05; **** p<0.0001.

Nitric oxide can be used for post-translational modification of proteins via S-nitrosylation (Stamler et al., 2001). Previous work has shown that increased nitric oxide has a suppressive role on lipolysis, and Auranofin, a thioredoxin reductase inhibitor that promotes S-nitrosylation, can inhibit lipolysis in 3T3-L1 cells (Yamada et al., 2015). Similarly, JS-K is a nitric oxide donor purported to promote S-nitrosylation, but it has not been shown to play a role in lipolysis (Nath et al., 2010; Shami et al., 2003). Given that both of these top hits were in the same pathway, we chose these for *in vivo* validation. We asked whether these drugs could modulate lipid droplet size and lead to increased adiposity in the zebrafish. We treated juvenile zebrafish for 24 hours with DMSO, Atglistatin, Auranofin, or JS-K and imaged the adipose tissue. We found that Atglistatin and JS-K significantly increase adipose tissue area and normalized area to standard length (Figure 5C, E). These effects were specific to the adipose tissue as standard length was not affected (Figure 5D). These data indicate that modulators of nitric oxide can inhibit lipolysis in cell lines, which then leads to an increase in adipose tissue area *in vivo* in the zebrafish. Moreover, this approach demonstrates the power of this system to dissect the relationship between novel modulators of lipolysis (i.e. nitric oxide) and adiposity *in vivo*.

### Lipolysis modulators also inhibit lipid droplet loss in melanoma cells

Upon uptake of adipocyte-derived lipids, cancer cells can store excess lipids in lipid droplets (Lengyel et al., 2018). Accumulation of lipid droplets in melanoma cells has been associated with increased metastatic potential and worse clinical outcomes (M. Fujimoto et al., 2020; Zhang et al., 2018). The mechanisms regulating subsequent lipolysis from the lipid droplets in cancer cells are not well understood, but we reasoned that some of the same mechanisms (i.e. ATGL, nitric oxide) used in adipocytes might also be used in cancer cells. To test this, we created a stable zebrafish melanoma cell line (ZMEL) that expressed the *ubb:plin2-tdtomato* construct (Heilmann et al., 2015) to generate the ZMEL-LD (lipid droplet) reporter cell line (Figure 6A). Because melanoma cells at baseline only have few small lipid droplets, we induced their formation via extrinsic addition of oleic acid, a key fatty acid that can be transferred from the adipocyte to the melanoma cell (Zhang et al., 2018). We found that after a pulse of oleic acid for 24 hours, we could easily detect PLIN2-tdTOMATO expression surrounding lipid droplets marked by the lipid droplet dye MDH (Figure 6B). A 3D reconstruction demonstrated that PLIN2-tdTOMATO was strictly expressed on the outline of the lipid droplet core, consistent with endogenous PLIN2 protein expression patterns (Olzmann & Carvalho, 2019) (Movie 1) and similar to what we saw in the adipocytes (Figure 1).

**Figure 6:**
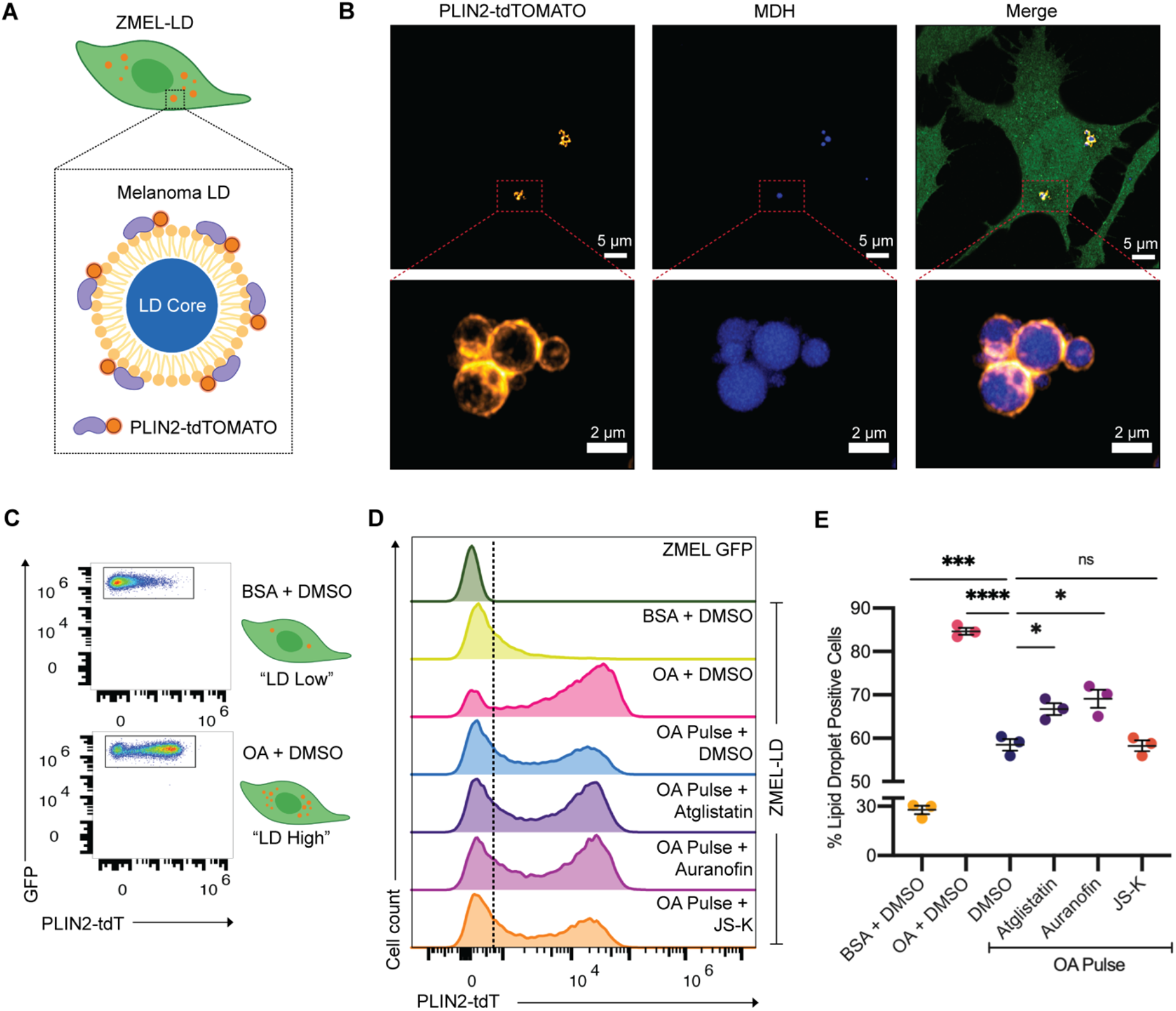
Lipolysis modulators also inhibit lipid droplet loss in melanoma cells. (A) Schematic of zebrafish melanoma cell line with lipid droplet reporter (ZMEL-LD) with lipid droplet labeled by PLIN2-tdTOMATO. (B) Confocal images of ZMEL-LD cells after 24 hours of oleic acid. Panels show fluorescence signal in PLIN2-tdTOMATO, MDH (lipid droplet dye) staining and merge of images with cytoplasmic GFP. Red box indicates position of higher magnification image of lipid droplets. (C) ZMEL-LD cells were treated with either BSA or oleic acid with DMSO for 72 hours then analyzed by FACS for PLIN2-tdTOMATO expression. (D) Representative histogram of PLIN2-tdTOMATO expression of ZMEL GFP and GFP+ ZMEL-LD cells with indicated drugs. Dashed line shows threshold for PLIN2-tdTOMATO expression. (E) Quantification of percent of GFP+ ZMEL-LD cells with lipid droplets. Lipid droplet low and high controls were ZMEL-LD cells treated with BSA or oleic acid for 72 hours. For drug treatments, ZMEL-LD cells were pulsed with oleic acid for 24 hours then given DMSO, 40 μM Atglistatin, 0.5 μM Auranofin, or 0.5 μM JS-K for 48 hours. N = 3 independent experiments. Bars indicate mean and SEM. Significance calculated via unpaired t-test; *p<0.05; ***p<0.001; ****p<0.0001.

To validate whether lipolysis inhibiting compounds could modulate lipid droplets in the ZMEL-LD cells, we utilized flow cytometry to measure PLIN2-tdTOMATO expression. We treated ZMEL-LD cells for 72 hours with either BSA or oleic acid as controls for low or high lipid droplet cell populations (Figure 6C), which confirmed the ability of the transgene to read out lipid droplets in this assay. We then tested the effects of the lipolysis inhibitors described above. We pulsed the ZMEL-LD cells with oleic acid for 24 hours (to induce lipid droplets) and then measured the subsequent decay in signal over the ensuing 48 hours, which is expected to decrease due to gradual lipolysis of the lipid droplets. Compared to cells with oleic acid pulse and DMSO (58.5 ± 1.4% LD+ cells), cells given JS-K (58.3 ± 1.2% LD+ cells) did not differ in the percent of lipid droplet positive cells (Figure 6D, E). In contrast, cells treated with Atglistatin (66.7 ± 1.4% LD+ cells) and Auranofin (69.1 ± 2.1% LD+ cells) demonstrated significantly higher lipid droplet positive cells (Figure 6D, E). These data indicate that similar to adipocytes, ATGL is a key regulatory step in lipolysis in the melanoma cells. Moreover, we find that nitric oxide, which was identified in our adipocyte screen, is similarly a modulator of lipolysis in the melanoma context and can be utilized for future studies to target adipocyte-melanoma cell cross-talk. We do not yet understand why different nitric oxide donors are more or less potent in adipocytes (where JS-K is a better inhibitor *in vivo*) versus melanoma cells (where Auranofin is a better inhibitor), but this could reflect differences in pharmacokinetics between the two cell types.

## Discussion

Lipid droplets are cytosolic storage organelles for cellular lipids which are dynamically regulated in response to metabolic and oxidative perturbations (Jarc & Petan, 2019). For instance, under hypoxic conditions, lipid droplets are crucial for protecting cells against reactive oxygen species and lipid peroxidation (Bailey et al., 2015; Bensaad et al., 2014). Lipid droplets can also buffer ER stress by sequestering excess lipids and proteins in the lipid droplet core (Chitraju et al., 2017; Velázquez et al., 2016; Vevea et al., 2015) while fluctuations in nutrient availability have been shown to lead to changes in lipid droplet biogenesis (Cabodevilla et al., 2013; Nguyen et al., 2017). The regulatory mechanisms driving these processes remain incompletely understood. Furthermore, lipid droplets are highly heterogeneous and the pathways which regulate lipid droplet dynamics in specific cell types warrant investigation.

To address such questions, we developed the first lipid droplet reporter in a vertebrate model organism. We show that our *plin2-tdtomato* reporter faithfully marks the lipid droplet *in vivo*. The combination of this reporter with the *in vivo* system of the *casper* zebrafish enables flexible and robust imaging approaches to examine lipid droplet regulation and function. In particular, the ease of chemical and genetic manipulation of the zebrafish combined with high-throughput imaging approaches enables interrogation of relevant pathways in a cell type specific manner. Furthermore, the capacity for intravital imaging creates the opportunity to conduct longitudinal analysis of lipid droplet dynamics across developmental time and in disease contexts between single animals.

Here, we demonstrate the capabilities of the tg(*ubb:plin2-tdtomato*) line by taking advantage of the fact that white adipocytes, which are primarily composed of a large unilocular lipid droplet (T. Fujimoto & Parton, 2011), are readily labeled by PLIN2-tdTOMATO expression. This labeling enables the study of individual adipocytes and adipose tissue in adult and juvenile zebrafish. We utilized this system to develop a robust imaging platform to specifically study the regulation of adipose tissue using both diet and pharmacologic perturbation. We focused on visceral adipose tissue due to its role as an endocrine organ and central regulator of organismal metabolism. Importantly, visceral adipose tissue accumulation, such as in obesity, influences the development of disorders including insulin resistance, cardiovascular disease, and hypertension (Fox et al., 2007; Le Jemtel et al., 2018; Verboven et al., 2018). We use our image analysis pipeline to demonstrate that our model is sensitive to diet induced changes in visceral adiposity. We also show that established chemical regulators of adipocyte lipolysis, Forskolin and Atglistatin, can produce quantitative changes in visceral adipose tissue. Collectively, these data illustrate the potential of our model to yield novel insights into the regulation of visceral adipose tissue, including in the context of obesity.

Although adipocytes comprise a major portion of adipose tissue, adipose tissue also consists of the stromal vascular fraction composed of fibroblasts, endothelial, and immune cells (Rosen & Spiegelman, 2014). Remodeling of adipose tissue architecture through changes in vascularization or recruitment of immune cells during tissue inflammation is associated with metabolic diseases including obesity and insulin resistance (Rosen & Spiegelman, 2014). Given the complexity of adipose tissue organization, understanding the native tissue architecture in relevant contexts is essential. We anticipate that our reporter can be easily crossed with other zebrafish transgenic reporters of interest to visualize heterotypic cell-cell interactions within adipose tissue.

While our studies show that this tool can be readily used to increase our understanding of adipocyte biology, it can also be utilized to study lipid droplets in other contexts as well. Lipid droplets are ubiquitous across almost all cell types. Therefore, this model could be applied to study the regulation of lipid droplets in the development and function of other adipose depots and additional cell types, such as muscle and hepatocytes (Bosma, 2016; Wang et al., 2013). In the disease context, we focused on the role of lipid droplets in cancer, since they have been implicated in various tumor types (Petan et al., 2018) where tumor cells can take up lipids from adipocytes and then package them into lipid droplets in the cancer cell (Balaban et al., 2017; Lengyel et al., 2018; Nieman et al., 2011; Zhang et al., 2018). This transfer of lipids has been linked to disease progression, making the regulation of lipid release from the lipid droplet through subsequent lipolysis in the tumor cell of particular interest. By expressing the *plin2-tdtomato* transgene in the ZMEL melanoma cells, we find that key regulators of lipolysis, such as ATGL and nitric oxide are mechanisms conserved with normal adipocytes. Interestingly, our results suggest that while the nitric oxide pathway can alter both adipose tissue area and lipid droplet content in melanoma cells, there may be differences between the phenotypes induced by nitric oxide production compared to more downstream effects such as S-nitrosylation, which are cell type specific. Collectively, this underscores the complexity of lipid droplet regulation and emphasizes the importance of studying these processes in both cell types. We believe that our model will serve as a powerful tool to study cell type specific regulation of lipid droplet biogenesis and function while preserving the endogenous structural and metabolic environment of an *in vivo* system.

## Acknowledgements

We thank members at the Memorial Sloan Kettering Cancer Center Aquatics Core, Molecular Cytology Core, and Flow Cytometry Core for their contributions to this work. We thank Dr. Mohita Tagore and Dr. Ting-Hsiang (Richard) Huang for comments on the project and manuscript.

## Competing Interests

D.L., E.J., J.W., E.M., O.O., and A.A. do not have competing interests to declare. R.M.W is a paid consultant to N-of-One Therapeutics, a subsidiary of Qiagen. R.M.W is on the scientific advisory board of Consano, but receives no income for this. R.M.W receives royalty payments for the use of the *casper* zebrafish line from Carolina Biologicals.

## Movie 1: 3D Reconstruction of ZMEL-LD Lipid Droplet

ZMEL-LD cells were given oleic acid for 24 hours, fixed and stained with the lipid droplet dye MDH. This movie is a 3D reconstruction of 37 planes covering a 6 μm stack of a lipid droplet cluster in a ZMEL-LD cell. PLIN2-tdTOMATO (orange) is located outside of the lipid droplet core (blue).

## Materials and Methods

### Cloning of ubb:plin2-tdtomato

To clone the *plin2* cDNA, tissue from the muscle and heart of adult *casper* zebrafish was dissected, pooled and then RNA was isolated using the Zymogen Quick RNA Miniprep Kit (Zymo Research, Irvine, USA, Catalog #R1054) according to manufacturer instructions. The Invitrogen SuperScriptIII First-Strand Synthesis SuperMix Kit (Thermo Fisher, Waltham, USA Catalog #18080400) was used according to manufacturer instructions to produce cDNA. CloneAmp HiFi PCR Premix (Takara, Mountain View, USA, Catalog #639298) was used to PCR amplify the PLIN2 cDNA and gel purified via NucleoSpin Gel and PCR Clean Up (Takara, Mountain View, USA, Catalog #740609.50). To generate pME-PLIN2-tdTOMATO, the PLIN2 cDNA was inserted on the 5’ end of pME-tdTOMATO using In-Fusion HD Cloning Plus (Takara, Mountain View, USA, Catalog #638920). Gateway cloning using the Gateway LR Clonase Enzyme mix (Thermo Fisher, Waltham, USA Catalog #11791019) was employed to create the *ubb:plin2-tdtomato* construct with p5E-ubb, pME-PLIN2-tdTOMATO, p3E-polyA into the pDestTol2pA2-blastocidin (cells) (Heilmann et al., 2015) or pDestTol2CG2 (zebrafish) (Kwan et al., 2007).

### Primers

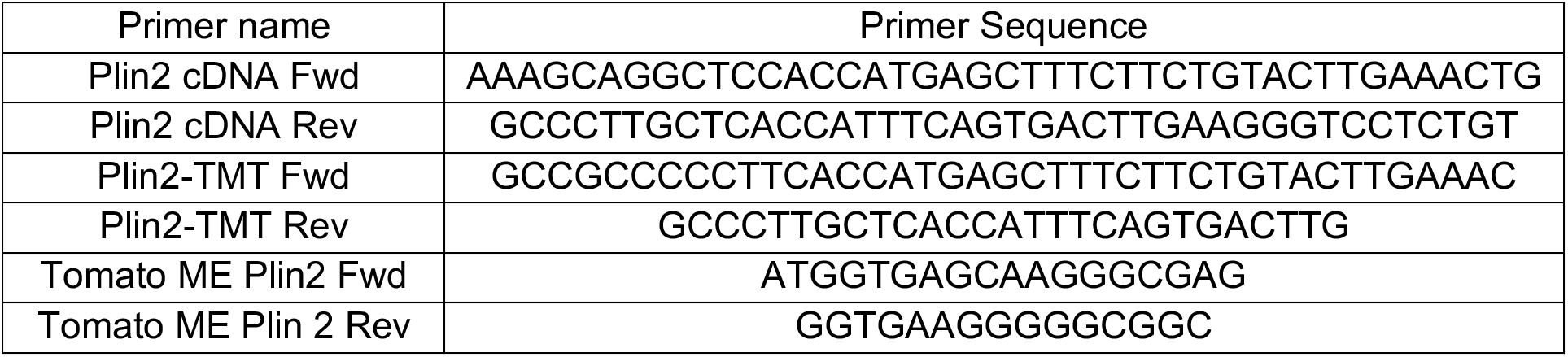

### Zebrafish Husbandry

All zebrafish experiments were carried out in accordance with institutional animal protocols. All zebrafish were housed in a temperature (28.5°C) and light-controlled (14 hours on, 10 hours off) room. Fish were initially housed at a density of 5 fish per liter and fed 3 times per day using rotifers and pelleted zebrafish food. All anesthesia was done using Tricaine (Western Chemical Incorporated, Ferndale, USA) with a stock of 4 g/L (protected for light) and diluted until the fish was immobilized. All procedures were approved by and adhered to Institutional Animal Care and Use Committee (IACUC) protocol #12-05-008 through Memorial Sloan Kettering Cancer Center.

### Generation of tg(ubb:plin2-tdtomato)

The *ubb:plin2-tdtomato* plasmid was injected into *casper* embryos with Tol2 mRNA to introduce stable integration of the *ubb:plin2-tdtomato* cassette. Fish with GFP+ hearts (due to pDestTol2CG) were selected and outcrossed to *casper* fish to produce the F1 generation. F1 zebrafish with GFP+ hearts and validated PLIN2-tdTOMATO expressing adipocytes were outcrossed to generate F2 generation zebrafish for experiments.

### Zebrafish Imaging and Analysis

Zebrafish were imaged using an upright Zeiss AxioZoom V16 Fluorescence Stereo Zoom Microscope with a 0.5x (for adult fish) or 1.0x (for juvenile fish) adjustable objective lens equipped with a motorized stage, brightfield, and GFP and tdTomato filter sets. To acquire images, zebrafish were lightly anesthetized with 0.2% Tricaine. Images were acquired with the Zeiss Zen Pro v2 and exported as CZI files for visualization using FIJI or analysis using FIJI (to manually quantify standard length) and MATLAB (Mathworks, Natick, USA).

Our adipocyte segmentation approach utilized the Image Processing Toolbox within MATLAB. Because the zebrafish gut is highly autofluorescent, we chose a threshold for the GFP channel to categorize as background signal and subtracted it from a determined threshold for the tdTOMATO channel. We used a set size to crop images around the tdTOMATO positive signal and created a mask for the adipose tissue. Within the masked area, we applied a higher tdTOMATO threshold to segment the fluorescent signal from the adipocytes. Finally, we quantified the number of pixels above the threshold to quantify adipose tissue area. For visualization purposes, the segmented images were color filtered on Adobe Photoshop from grayscale to gold color scale.

### BODIPY staining of zebrafish

Adult zebrafish were placed in tanks and juvenile zebrafish were placed in p1000 tip boxes with either DMSO or 10 ng/μL BODIPY 493/503 (Thermo Fisher, Waltham, USA, Catalog #D3922) for 30 mins in the dark. Fish were washed then placed in new tanks with fresh water for 1 hour. Fish were washed again to remove any residual BODIPY then anesthetized and imaged as indicated above for whole adipose tissue.

Higher resolution images of zebrafish adipocytes were acquired using the Zeiss LSM 880 inverted confocal microscope with using a 10x objective. Zebrafish were lightly anesthetized with 0.2% Tricaine and mounted on a glass bottom dish (MatTek, Ashland, USA, Catalog #P35G-1.5-20-C) with 0.1% low gelling agarose (Sigma-Aldrich, St. Louis, USA, Catalog, #A9045-25G).

### IHC for tdTOMATO

Zebrafish were sacrificed in an ice bath for at least 15 minutes. For adults, zebrafish tails were dissected. For juvenile zebrafish, the entire fish was used for fixation. Selected zebrafish were fixed in 4% paraformaldehyde for 72 hours at 4°C, washed in 70% ethanol for 24 hours, and then paraffin embedded. Fish were sectioned at 5 μm and placed on Apex Adhesive slides, baked at 60°C, and then stained with antibodies against tdTomato (1:500, Rockland, #600-401-379). All histology was performed and stained by Histowiz.

### Juvenile Zebrafish Fast

tg(*ubb:plin2-tdtomato*) F1 fish were outcrossed to *caspers* to generate the F2 generation. F2 fish were raised at a standard density of 25 fish per 2.8 L tank. At 21 dpf, fish were separated into new tanks which received standard feed or were fasted for 2.5 days. Fish were anesthetized with tricaine and imaged as described above to quantify visceral adipose tissue area and standard length.

### High-fat diet feeding

tg(*ubb:plin2-tdtomato*) F2 zebrafish were raised at a standard density of 25 fish per 2.8 L tank. At 21 dpf, the zebrafish were placed into 0.8L tanks and fed either a high fat or control diet (Sparos, Portugal) for 7 days. Fish were then imaged for Plin2-tdtomato expression at 28 dpf. Prior to imaging, fish put in a new tank and food withheld for ~16-20 hours. Zebrafish were at equal density for control and experimental groups, ranging from 15-30 fish per tank. Fish were fed 0.1 g feed per tank per day split over two feedings. The high fat and control diets were customized and produced at Sparos Lda (Olhão, Portugal), where powder ingredients were initially mixed according to each target formulation in a double-helix mixer, being thereafter ground twice in a micropulverizer hammer mill (SH1, Hosokawa-Alpine, Germany). The oil fraction of the formulation was subsequently added and diets were humidified and agglomerated through low-shear extrusion (Dominioni Group, Italy). Upon extrusion, diets were dried in a convection oven (OP 750-UF, LTE Scientifics, United Kingdom) for 4 h at 60 °C, being subsequently crumbled (Neuero Farm, Germany) and sieved to 400 microns. Experimental diets were analyzed for proximal composition. The Sparos control diet contains 30% fishmeal, 33% squid meal, 5% fish gelatin, 5.5% wheat gluten, 12% cellulose, 2.5% Soybean oil, 2.5% rapeseed oil, 2% vitamins and minerals, 0.1% vitamin E, 0.4% antioxidant, 2% monocalcium phosphate, and 2.2% calcium silicate. The Sparos HFD contains 30% fishmeal, 33% squid meal, 5% fish gelatin, 5.5% wheat gluten, 12% palm oil, 2.5% soybean oil, 2.5% rapeseed oil, 2% vitamins and minerals, 0.1% vitamin E, 0.4% antioxidant, 2% monocalcium phosphate, and 2.2% calcium silicate.

### 3T3-L1 Cell Culture

3T3-L1 cells were acquired from ZenBio and followed their differentiation protocol. Cells are received at Passage 8 and split to a maximum of Passage 12 as per recommendation of the company. 96-well plates were coated with fibronectin (EMD Millipore, Burlington, USA, Catalog #FC010) diluted 1:100 in PBS for at least 30 minutes to promote improved adherence of cells to the dish. 3T3-L1 cells are first cultured in PM-1-L1 Preadipocyte Medium and allowed to grow to 100% confluence. PM-1-L1 media is changed every 48-72 hours. 48 hours after reaching 100% confluence, cells were changed to DM-1-L1 Differentiation Medium for 72 hours and then changed to AM-1-L1 Adipocyte Media. AM-1-L1 Adipocyte Media was changed every 48-72 hours. Once in AM-1-L1, media is changed gently with a multichannel pipette and only 150μL of the 200μL is replaced to prevent touching the bottom of the well with the pipette tip. After 2-3 weeks in AM-1-L1, the 3T3-L1 develop significantly large lipid droplets and were used in the screen.

### LOPAC Library Screen

The LOPAC library includes 1280 clinically relevant compounds with annotated targets or pathways. The workflow of the screen involved drug or vehicle control of the 3T3-L1 adipocytes for 24hrs in serum free media. After 24 hours, 100 μL of the media supernatant was collected to measure secreted glycerol using the Free Glycerol Reagent (Sigma-Aldrich, St. Louis, USA, Catalog F6428) and following the associated glycerol assay protocol.

Media (screen media) used for drug treatment was phenol-free DMEM supplemented with 0.2% BSA FFA-free (Sigma-Aldrich, St. Louis, USA, Catalog 9048-46-8). The 1280 compounds were aliquoted as 2 μL at 1 mM into 16x 96-well plates and stored at −20C. Upon thawing, 198 μL of screen media was added to the well, bringing the final drug concentration for all compounds in the screen to 10 μM. Control vehicle was 1% DMSO served as a negative control and 1uM Isoproterenol served as a positive control in the screen. This media containing LOPAC drugs, DMSO, and Isoproterenol was transferred to 3T3-L1 cells and incubated for 24 hours.

To measure glycerol release as a readout for lipolysis, 100 μL of Free Glycerol Reagent was aliquoted per well of a 96 well plate. 10 μL of supernatant media from 3T3-L1 adipocytes was then added to each well. A standard curve was produced by using Glycerol Standard Solution (Sigma-Aldrich, St. Louis, USA, Catalog G7793). The plate is incubated at 37C for 5 minutes and then developed with a plate reader set to detect absorbance at 540 nm. Using the standard curve, a fit equation is developed in Excel to convert the absorbance values into glycerol concentration. To take into account differences that occur in wells on the edge versus middle of the plate, all well positions across all plates in the screen are averaged to create a normalization factor for any given position on the plate. These normalized values were then used to determine top hits for compounds either that block lipolysis.

### Glycerol Release Assay with Atglistatin

3T3-L1s were differentiated on a fibronectin-coated 96-well dish. At the start of the lipolysis experiment, 3T3-L1s were changed to serum-free DMEM supplemented with 0.2% BSA FFA-free (Sigma-Aldrich, St. Louis, USA, Catalog 9048-46-8). The media was supplemented with 1% DMSO for negative control or 1uM isoproterenol to induce lipolysis +/- 100 μM Atglistatin (Sigma-Aldrich, St. Louis, USA, Catalog SML1075) to block lipolysis and cells were incubated for 24 hours.

### To measure glycerol release, 100 μL of Free Glycerol Reagent was aliquoted per well of a new

96 well plate. 10 μL of supernatant media from 3T3-L1 adipocytes was then added to each well. A standard curve was produced by using Glycerol Standard Solution (Sigma-Aldrich, St. Louis, USA, Catalog G7793). The plate is incubated at 37C for 5 minutes and then developed with a plate reader set to detect absorbance at 540 nm. Using the standard curve, a fit equation is developed in Excel to convert the absorbance values into glycerol concentration.

### Juvenile Zebrafish Drug Treatments

*tg*(*ubb:plin2-tdtomato*) F1 fish were outcrossed to *caspers* to generate the F2 generation. F2 fish were raised at a standard density of 50 fish per 6.0L tank. For drug treatment, fish were removed from the system at 21 dpf and placed at a density of 1 fish per well in a 6 well plate with 10 mL of E3 per well. After a 24 hour incubation with the drug fish were anesthetized with Tricaine and imaged using the described protocol to quantify (1) Standard length and (2) area of PLIN2-tdTOMATO expression corresponding to visceral adipose tissue area. Fish were treated with the following compounds, which were all dissolved in DMSO.

| Compound | Dose | Source |
| --- | --- | --- |
| Forskolin | 5 μM | Sigma-Aldrich, St. Louis, USA, Catalog, Catalog #F6886 |
| Auranofin | 1μM | Sigma-Aldrich, St. Louis, USA, Catalog, Catalog #A6733 |
| JS-K | 1 μM | Sigma-Aldrich, St. Louis, USA, Catalog, Catalog #J4137 |
| Atglistatin | 40 μM | Sigma-Aldrich, St. Louis, USA, Catalog, Catalog #SML1075 |

### Generation of ZMEL-LD Cell Line

The ZMEL zebrafish melanoma cell line was derived from a tumor of a *mitfa:BRAF^V600E^/p53^-/-^* zebrafish was described previously (Heilmann et al., 2015). ZMEL cells constitutively express eGFP driven by the *mitfa* promoter(Heilmann et al., 2015). ZMEL cells were grown at 28°C in a humidified incubator in DMEM (Gibco, Waltham, USA, Catalog, #11965) supplemented with 10% FBS (Gemini Bio, #100-500), 1X penicillin/streptomycin/glutamine (Gibco, Waltham, USA, Catalog, #10378016), and 1X GlutaMAX (Gibco, Waltham, USA, Catalog, #35050061). To generate the ZMEL-LD cells, ZMEL cells were nucleofected with the *ubb:plin2-tdtomato* plasmid using the Neon Transfection System (Thermo Fisher, Waltham, USA, Catalog #MPK10096), selected for two weeks in blasticidine supplemented media at 4 μg/μL (Sigma-Aldrich, St. Louis, USA, Catalog, #15205-25MG), and FACS sorted for GFP and tdTOMATO double positive cells.

### ZMEL-LD Imaging

8 well Nunc Lab-Tek Chambered Coverglass was coated with 1:100 dilution of fibronectin in DPBS (Millipore Sigma, Burlington, USA, Catalog #FC010-5MG) for 30 mins and then washed with DPBS (Thermo Scientific, Waltham, USA, Catalog, #14190-250). ZMEL-LD cells were seeded at 30,000 cells per well and left to adhere for 24 hours. Media supplemented with 250 μM oleic acid (Sigma-Aldrich, St. Louis, USA, Catalog, #O3008-5ML) was added for 24 hours. Cells were fixed with 2% paraformaldehyde (Santa Cruz Biotechnology, Santa Cruz, USA, Catalog #sc-281692) for 45 minutes, washed with DPBS and permeabilized with 0.1% triton-X (Thermo Fisher, Waltham, USA, Catalog #PI85111) for 30 minutes at room temperature. To stain for lipid droplets, cells were washed and stained with 1:500 MDH (Abcepta, San Diego, USA, Catalog #SM1000a) for 15 minutes. Cells were imaged on the Zeiss LSM 880 inverted confocal microscope with AiryScan using a 63x oil immersion objective. Confocal stacks were visualized via FIJI and 3D reconstruction was created using Imaris (Bitplane Inc, Concord, USA).

### ZMEL-LD FACS Analysis

ZMEL Dark (no fluorescence), ZMEL-GFP, ZMEL-LD cells were plated on fibronectin coated 6 well plates at a density of 500,000 cells in 1 mL of media per well. At 24 hours after plating, cells were given either 150 μM of BSA or oleic acid with 1 μL of DMSO. At 48 and 72 hours after plating, lipid droplet low and high controls were switched to fresh media with 150 μM of BSA or oleic acid with 1 μL of DMSO. Cells pulsed with oleic acid received fresh media with 150 150 μM of BSA with either 40 μM Atglistatin, 0.5 μM Auranofin or 0.5 μM JS-K. At 96 hours after plating, cells were trypsinized, washed with DPBS and resuspended in DMEM supplemented with 2% FBS, 1X penicillin/streptomycin/glutamine, and 1X GlutaMAX. Cells were stained for viability with 1:1000 DAPI and strained through the Falcon FACS Tube with Cell Strainer Cap (Thermo Fisher, Waltham, USA Catalog, #08-771-23). Data was acquired via the Beckman Coulter CytoFLEX Flow Cytometer (Beckman Coulter, Miami, USA) and analyzed via FlowJo software (BD Biosciences, San Jose, USA).

### Schematics

Schematics and illustrations were generated via Biorender on biorender.com.

### Statistics

All statistical analysis was performed using GraphPad Prism 8 (Graphpad, San Diego, USA). Data are presented as mean ± standard error (SEM). P < 0.05 was considered statistically significant. Statistical tests used were Mann-Whitney, Kruskal-Wallis or unpaired t-tests which are noted in the figure legend. All experiments were done with at least 3 independent replicates. For *in vivo* experiments, N denotes number of independent experiments while n denotes number of individual fish. Imaging analysis utilized FIJI, Imaris, MATLAB software.

